## Supplemental Figures for "Monoclonal antibodies derived from B cells in subjects with cystic fibrosis reduce *Pseudomonas aeruginosa* burden in mice"

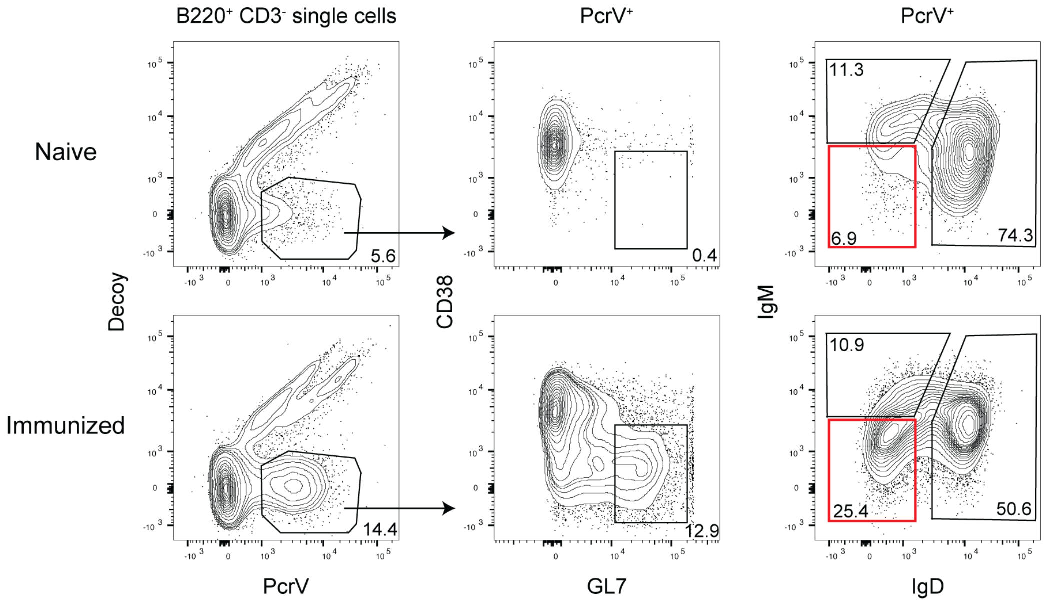


**Figure 1–figure supplement 1. Tetramer-specific class-switched B cells in mice immunized with PcrV.** Flow cytometry plots from lymphoid tissue in representative PcrV-immunized or control (naïve) mice sacrificed on day 7 post-immunization by intraperitoneal injection. Cells were analyzed after the magnetic enrichment of tetramer-bound cells. Class-switched B cells are highlighted within the red boxes.

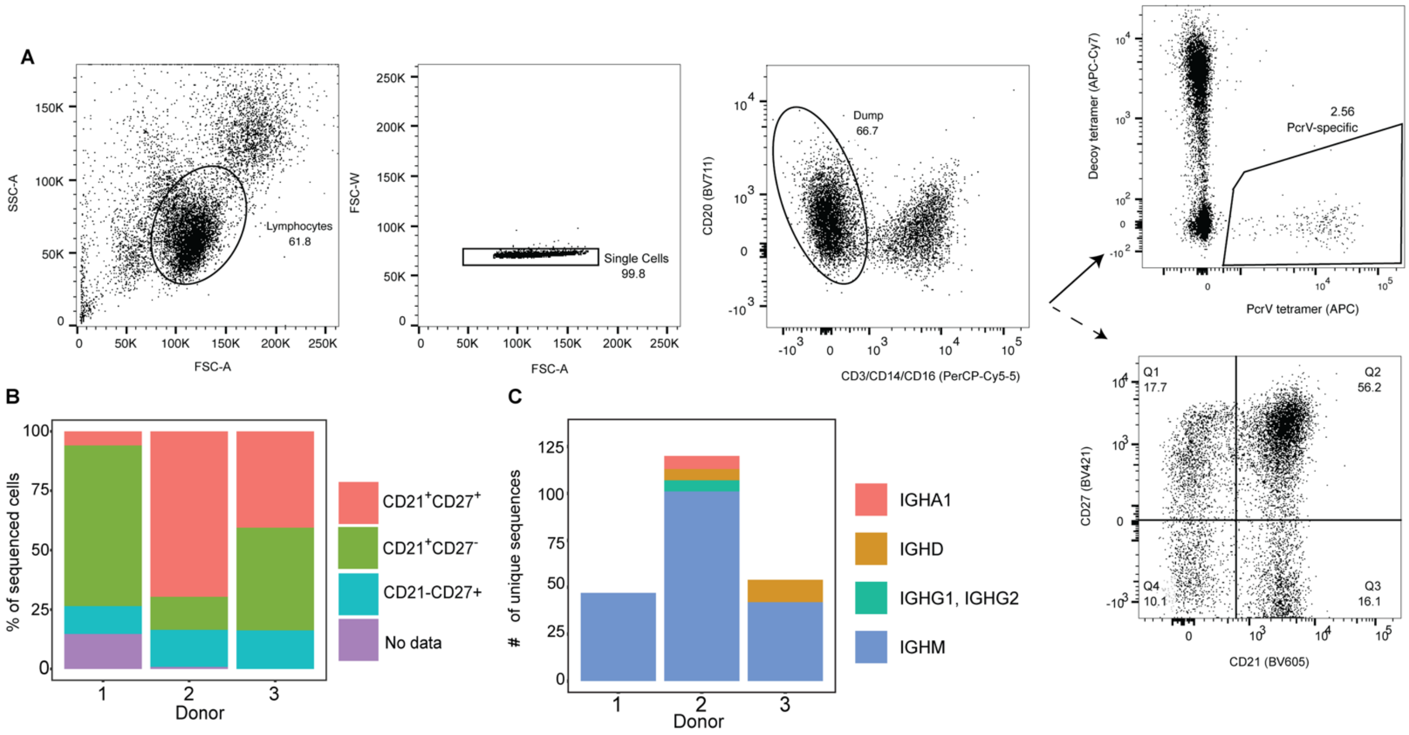


**Figure 2–figure supplement 2. Summary of data generated by single cell BCR sequencing.** A) Flow cytometry gating strategy. B) Surface marker expression of singly-sorted cells used for BCR sequencing for each donor (x-axis). C) Enumeration of unique heavy chain sequences obtained from PcrV-specific B cells in each donor, color-coded by the sequenced constant region.

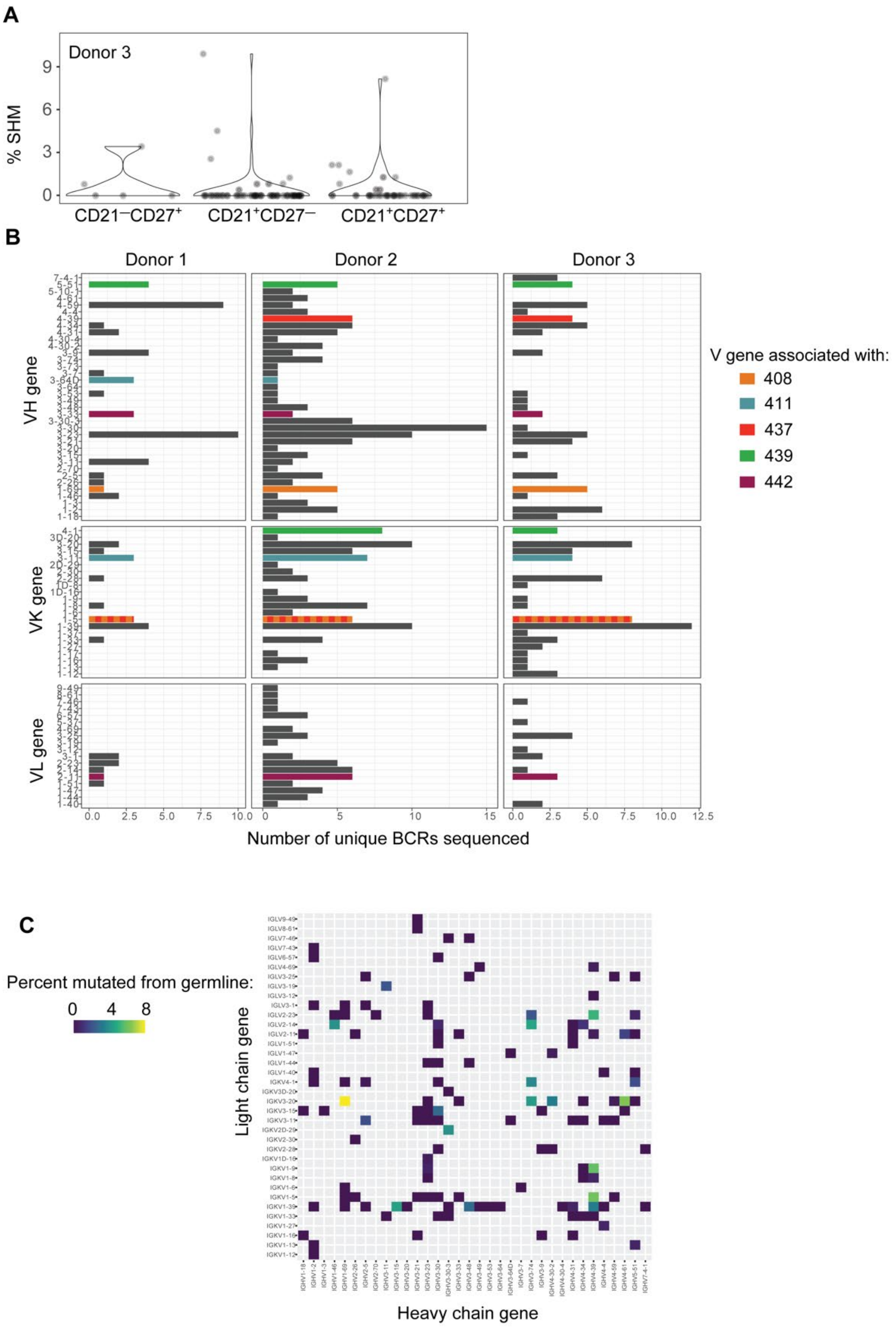


**Figure 4–figure supplement 3**. **B cell receptor (BCR) sequencing of PcrV-tetramer-specific B cells derived from 3 CF donors.** A) Percentage of somatic hypermutation (SHM) detected in BCR sequences from cells of the indicated phenotype in CF donor 3. Each circle represents a heavy or light chain sequence from a singly-sorted cell. B) Histograms show the number of

unique BCR sequences obtained for each V gene (y-axis) for heavy and light (kappa or lambda) chains. Data from each CF donor (Donors 1-3) are shown in a separate panel of graphs (with Donor number indicated at top). The bars for the V genes used by *in vivo*-tested mAbs are colored as indicated. C) Heatmap showing pairings of heavy- (x-axis) and light (y-axis) chain V genes for 175 BCR sequences where full-length, high quality V region sequences were attained. For each heavy/light chain pair, the percentage of heavy chain sequence which differs from the germline sequence is depicted by color gradient.

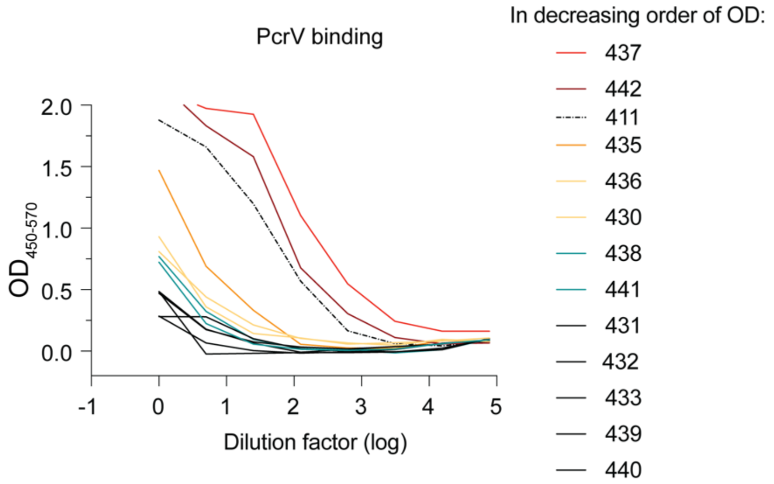


**Figure 4–figure supplement 4**. **Transfectant supernatant screen of 12 MBC-derived mAbs.** ELISA assessing PcrV binding for supernatants from 293T cells transfected with IgG expression plasmids. Numbers (430-442) indicate the BCR sequences identified from individual, PcrV-specific, MBCs derived from CF donor 2. Supernatant for mAb 411 IgG (dotted line; isolated from CF Donor 1) is included as a benchmark.
